# Factors that affected the qualification of standardized residency training of internal medicine: a prospective longitudinal study

**DOI:** 10.1101/2022.02.15.480480

**Authors:** Yunjuan Sun, Jialiang Xu, Limei Hu, Jiajun Qi

**Author notes:** **Corresponding Author: Jiajun Qi**, M.D., Education Training Center, the First Affiliated Hospital of Soochow University. No. 899 Pinghai Road, Gusu District, Suzhou City, Jiangsu Province, the People’s Republic of China, 215006. Contributions: Conceptualization, Formal analysis, Investigation and Writing original draft. Contributions: Methodology, Investigation, Review and editing. Contributions: Collection of data and Supervision. Contributions: Conceptualization, Methodology, Investigation, Supervision, Review and editing.

## Abstract

**Introduction:** Since the standardized residency training (SRT) was launched in 2014, there were very few data on evaluation of the factors that could influence the residents’ competency in China.

**Methods:** All residents who started their SRT programme from September 2015 to September 2018 were enrolled in this study. During the following years, they finished the programme and took examinations. The demographic information and exam scores of each subject were collected and analyzed.

**Results:** We found that the qualification rates of SRT programme differed significantly annually. Age, gender, the highest medical education degree, residents from different places, training duration and years of medical school study were also different significantly year by year. According to the scores of examinations, residents were divided into two groups: the qualified group and the non-qualified group. Age, the highest medical education degree, residents from different places, training duration, years of medical school study and prior medical work experience differed significantly between the two groups. Multivariate Logistic regression analysis showed the residents from different places and the training duration were factors that affected the qualification rate of SRT programme independently (*P* = 0.000 and 0.005, respectively).

**Discussion:** The “homogeneity” of SRT had been achieved regardless of the individual discrepancy, while the resident from different places and the training duration were obstacles we need to overcome further. Therefore, we should further advance the process of the transition from “Unit Persons” to “Society Persons”. And extending training duration will facilitate their qualification and be benefit to improve their competency.

## Introduction

China is home to nearly one fifth of the world’s population while the shortage of medical care personnel is a long-standing problem. In 2010, the number of doctors per 1000 population was 1.43 in China which was almost half of the average level of United States and United Kingdom [1]. The discrepancy in the level of medical education and competence of Chinese doctors is great. In China, the medical education qualification certificates include secondary vocational diploma (Zhong Zhuan), vocational diploma (Zhuan Ke), bachelor’s degree, master’s degree and doctoral degree. After graduating from the health vocational school or medical school, they could work under the supervision of qualified medical practitioners. Before 2014, they were not required to take residency training in recognized specialties except one year internship in health work unit. Then, they became assistant doctors or doctors after passing the National Medical Licensing Examination. Furthermore, for nearly half a century, our medical schools have trained the students without distinguishing degree types between professional and academic degrees. Because the relatively short period of medical training time, the academic degree graduates were generally thought to be with relatively weak clinical capability and difficult to meet the clinical needs. Thus, the shortage of enough quantity and quality qualified doctors, especially physicians, severely restricted the improvement of basic medical and health services [2]. This dilemma catalyzed a health care reform of resident training in a top-down manner of the country. In Dec 2013, the guidance on the establishment of standardized training system for residents was promulgated by the National Health Commission of the People’s Republic of China and other seven departments, finally [3]. Following the pilot model developed in Shanghai, called 5+3 [4], the standardized residency training (SRT) programme would be launched comprehensively among all the provinces by 2015. All medical clinicians with bachelor’s degree and above would receive standardized training for residents [3].

Now, SRT is a mandatory residency training for medical graduates before they become independent healthcare practitioners. Based on their different educational background, graduates with bachelor’s, master’s or doctor’s degree are asked to receive the SRT for three-year two-year and one-year, respectively. This programme contains 34 subspecialties, covering almost all aspects of clinician education. In the SRT programme, residents are no longer fixed in one post, instead, they will rotate among different departments. After they accomplished the programme, they could apply for a stable position. As internal medicine residency training, residents with bachelor’s degree are required to carry on 33 months training in different departments of internal medicine, including cardiology, pulmonology, gastroenterology, haematology, nephrology, rheumatology, endocrinology, emergency medicine, critical care medicine, psychology counselling clinic, outpatient and other elective department. Residents will strengthen their clinical competencies on rotation which was consisted of professional skills, theoretical knowledge, and communication techniques aspects. According to the guidelines, the assessments of residents are conducted at each department and annually. All the residents must have accomplished their entire training plan. Then, they could have the chance to apply for the exit examination, which tests the theoretical knowledge and clinical skills of residents. Those who failed to pass the standardized exit examination would not receive a certification of the programme and could not become independent practitioners in local area.

And, teaching hospitals affiliated with universities are approved to carry out the SRT programme according to the Standards of Training Bases released by the National Health Commission. The training bases involve three types of hospitals, namely, tertiary general, tertiary specialty, and secondary general hospitals. These bases are supported by the National Health Commission, the State Commission Office for Public Sector Reform, the National Development and Reform Commission, the Ministry of Education, the Ministry of Finance, the Ministry of Human Resources and Social Security and the State Administration of Traditional Chinese Medicine as well as their subordinate units and departments. Furthermore, the expert committees from the Chinese Medical Association hold training conferences every year to train the trainers which guaranteed the high-quality and standardization of training. In addition, the SRT programme integrated with the medical professional degree postgraduate education (eg, MM and MD). Postgraduates who passed the national entrance examination for postgraduate at the beginning of SRT could simultaneously obtain a certificate of residency training and a certificate of professional degree of postgraduate education at the end of the programme [5,6]. Thus, the trainees are comprised of the residents who graduated from medical schools and first-year postgraduates who just enrolled in medical professional degree postgraduate education.

The original intention of SRT is to achieve homogeneous competency among residents. Yao He and et, al. found that SRT have achieved a standardized training quality across different types of teaching hospitals despite of residents’ individual demographic background and contextual factors [7]. However, there are still a few residents could not pass the exit examination and failed to graduate from the programme. There are very few data on evaluation of SRT in China addressing on this topic. To provide more evidence, the present study focused on the factors that might affect the qualification of SRT programme of internal medicine.

## Methods

### Participants

All 446 residents who started their internal medicine residency training from September 2015 to September 2018 in our training base were enrolled. They came from 13 hospitals which located in the Suzhou Prefecture, Jiangsu Province. This training base was a tertiary general hospital affiliated to the Soochow University (SUDAH). The highest educational degree of each resident was bachelor’s, master’s or doctoral degrees. According to the different places where the residents came from, trainees were divided into three groups: SUDAH residents, local hospital residents and clinical postgraduates. The SUDAH residents were the residents who were newly recruited by our hospital as new staffs. The local hospital residents were the residents who were referred to our base for the STR programme from the secondary general or specialty hospitals. These secondary hospitals had signed contracts of employment with the local resident trainees in advanced. The clinical postgraduates were the students who continued their postgraduate studies in Soochow University after they graduated from 5-year medical schools. According to the guidelines of SRT, residents were differed in training duration based on their highest medical education degrees: one-, two- and three-year. For the residents with doctoral degree, the training duration would be three-year for the residents with academic degree and one-year for the residents with professional degree, respectively. For the residents with master’s degree, the training duration could be three-year for the residents with academic degree and two-year for the residents with professional degree, respectively. The training duration for bachelor’s degree was three-year. Therefore, the residents entered the programme in the same year might be exited in different years. The last session of trainees in the present study were enrolled in September 2018 and took the exit examinations in February 2020. The years of medical school study were the years spending on studying for medical educational degrees.

### Program description

The demographic information of each subject was collected, including age, gender, the highest medical education degree and the places where the residents came from (SUDAH residents, local hospital residents and clinical postgraduates). The information of SRT programme, including training duration, years of medical school study, prior medical work experience. The first and second examination scores and final qualification rate were also gathered. All the residents would be assessed twice except the residents with one-year training duration, who only took examination once. The examination consisted of two parts: written and hands-on part. The full mark of each part was 100 points and the acceptance line was 60 points. The qualification meant the scores of both parts would meet with the criteria. The work was carried out in accordance with the Declaration of Helsinki. Ethical approval has been granted by the First Affiliated Hospital of Soochow University Institutional Review Board.

### Data analysis

Continuous variables were expressed as the mean ± standard deviation, and categorical variables were expressed as the number and percentage. A Student *t*-test or Mann-Whitney U test was used, as appropriate, to analyze the differences in the continuous variables, and the chi-square test was used to analyze the differences in the dichotomous variables unless the expected values in the cells were < 5, in which case a Fisher’s exact test was used. All variables with a *P* < 0.05 were then tested in a forward stepwise multivariate Logistic regression analysis. Data were expressed as odds ratio with 95% confidence interval. *P* < 0.05 was considered statistically significant. All statistical analyses described above were performed with SPSS version 26 software (IBM SPSS Statistic Institute, Chicago, US).

## Results

### The demographic data and qualification rate of residents each year

From 2015 to 2018, the demographic data and the examination scores of all the residents were shown in Table 1 by the year. Age, gender, the highest medical education degree, residents from different places, training duration, years of medical school study, examination scores and qualification rates differed significantly every year, except the prior medical work experience. Figure 1 illustrated the gender distribution of the residents by the year (*P* = 0.040, supplements). The residents with bachelor’s degree accounted for a growing proportion by the year (Figure 2A, supplements). The percentages of residents training duration of three years increased annually (*P* = 0.001, Figure 2B, supplements). Each examination score was illustrated annually in Figure 3 (all of *Ps* < 0.001, supplements).

**Table 1.** Comparisons of the demographic data and the examination performances annually.

| Variate | 2015 | 2016 | 2017 | 2018 | *P* value |
| --- | --- | --- | --- | --- | --- |
| Age (yr.) | 25.77±2.45 | 25.75±2.42 | 24.88±2.12 | 24.43±2.03 | < 0.001 |
| Male (%) | 40 (35.7) | 37 (25.9) | 18 (18.2) | 27 (29.3) | 0.040 |
| Highest medical education degree |  |  |  |  | 0.010 |
| Bachelor's (%) | 76 (67.9) | 88 (61.5) | 76 (76.8) | 80 (87.0) |  |
| Master's (%) | 27 (24.1) | 44 (30.8) | 18 (18.2) | 7 (7.6) |  |
| Doctoral (%) | 9 (8.0) | 11 (7.7) | 5 (5.1) | 5 (5.4) |  |
| Residents from different places |  |  |  |  | < 0.001 |
| SUDAH residents (%) | 34 (30.4) | 42 (29.4) | 16 (16.2) | 10 (10.9) |  |
| Clinical postgraduates (%) | 70 (62.5) | 71 (49.7) | 71 (71.7) | 77 (83.7) |  |
| Local hospital residents (%) | 8 (7.1) | 30 (21.0) | 12 (12.1) | 5 (5.4) |  |
| Training duration |  |  |  |  | 0.001 |
| One-year (%) | 5 (4.5) | 2 (1.4) | 5 (5.1) | 0 (0) |  |
| Two-year (%) | 20 (17.9) | 31 (21.7) | 12 (12.1) | 3 (3.3) |  |
| Three-year (%) | 87 (77.7) | 110 (76.9) | 82 (82.8) | 89 (96.7) |  |
| Years of medical school study (yr.) | 6.04±1.65 | 6.24±1.71 | 5.82±1.63 | 5.51±1.42 | 0.006 |
| Prior medical work experience (%) | 18 (16.1) | 30 (21.0) | 11 (11.1) | 13 (14.1) | 0.200 |
| Fist written exam score (points) | 64.11±10.63 | 72.21±8.95 | 70.77±7.48 | 69.25±9.26 | < 0.001 |
| Fist hands-on exam score (points) | 90.15±2.72 | 90.29±3.79 | 87.15±4.36 | 82.96±4.57 | < 0.001 |
| Second written exam score (points) | 66.57±6.29 | 64.07±6.26 | 77.78±8.07 | 75.61±8.39 | < 0.001 |
| Second hands-on exam score (points) | 90.52±3.51 | 86.47±9.27 | 79.82±4.62 | 84.31±7.83 | < 0.001 |
| Qualification (%) | 77(68.8) | 124(86.7) | 91(91.9) | 87(94.6) | 0.000 |

### Comparisons between the qualified and the non-qualified groups

According to the acceptance lines of criteria set priorly, residents were divided into two groups: the qualified group and the non-qualified group. The comparisons of the demographic data and the exam scores between the two groups were shown in Table 2. Residents of the non-qualified group was older than the qualified group (26.13±2.46 vs 25.14±2.30, *P* = 0.001). And, distributions of the highest medical education degree of the two groups differed significantly (*P* = 0.006). The percentage of doctoral degree of the non-qualified group was higher than the qualified group (16.4% vs 5.0%) and the percentage of bachelor’s degree of the former group was lower than the latter group (58.2% vs 74.1%). Therefore, the years of medical school study of the non-qualified group was a little longer than the other group (6.58±2.08 vs 5.84±1.53, *P* = 0.006). More residents in the non-qualified group had prior medical work experience than the qualified group (26.9% vs 14.2%, *P* = 0.010). Figure 4A illustrated the differences of percentages of the residents came from different places (*P* = 0.000, supplements). In the non-qualified group, the percentage of clinical postgraduates was obviously lower than the qualified group (41.8% vs 68.9%). Also, there was a difference in the training duration between the two groups (*P* = 0.002, Figure 4B, supplements). The percentage of one-year training duration in the non-qualified group was obviously higher than the qualified group (10.4% vs 1.3%).

**Table 2.** Comparisons of the demographic data and examination performances between the qualified and non-qualified groups.

| Variate | Qualified | Non-qualified | *P* |
| --- | --- | --- | --- |
| Age (yr.) | 25.14±2.30 | 26.13±2.46 | 0.001 |
| Male (%) | 103(27.2) | 19(28.4) | 0.841 |
| Highest medical education degree |  |  | 0.004 |
| Bachelor's (%) | 281(74.1) | 39(58.2) |  |
| Master's (%) | 79(20.8) | 17(25.4) |  |
| Doctoral (%) | 19(5.0) | 11(16.4) |  |
| Residents from different places |  |  | 0.000 |
| SUDAH residents (%) | 78(20.6) | 24(35.8) |  |
| Clinical postgraduates (%) | 261(68.9) | 28(41.8) |  |
| Local hospital residents (%) | 40(10.6) | 15(22.4) |  |
| Training duration |  |  | 0.002 |
| One-year (%) | 5(1.3) | 7(10.4) |  |
| Two-year (%) | 56(14.8) | 10(14.9) |  |
| Three-year (%) | 318(83.9) | 50(74.6) |  |
| Years of medical school study (yr.) | 5.84±1.53 | 6.58±2.08 | 0.006 |
| Prior medical work experience (%) | 54(14.2) | 18(26.9) | 0.010 |
| Fist written exam score (points) | 70.96±8.11 | 57.28±7.92 | 0 |
| Fist hands-on exam score (points) | 88.04±5.07 | 87.82±4.70 | 0.760 |
| Second written exam score (points) | 71.82±8.51 | 62.20±9.90 | 0 |
| Second hands-on exam score (points) | 84.24±6.63 | 84.72±13.17 | 0.803 |

### Factors that affected the qualification rate

Multivariate Logistic regression analysis showed that resident from different places and training duration were independently associated with qualification (*P* = 0.000 and *P* = 0.005, respectively). Confidence intervals and significance levels were presented in Table 3. SUDAH residents and clinical postgraduates were favorable factors for qualification (OR= 0.614 and 0.208). The training duration for residents with two-year and three-year were easier to pass the STR programme than residents with one-year training duration (OR=0.104 and 0.226, respectively).

**Table 3.**
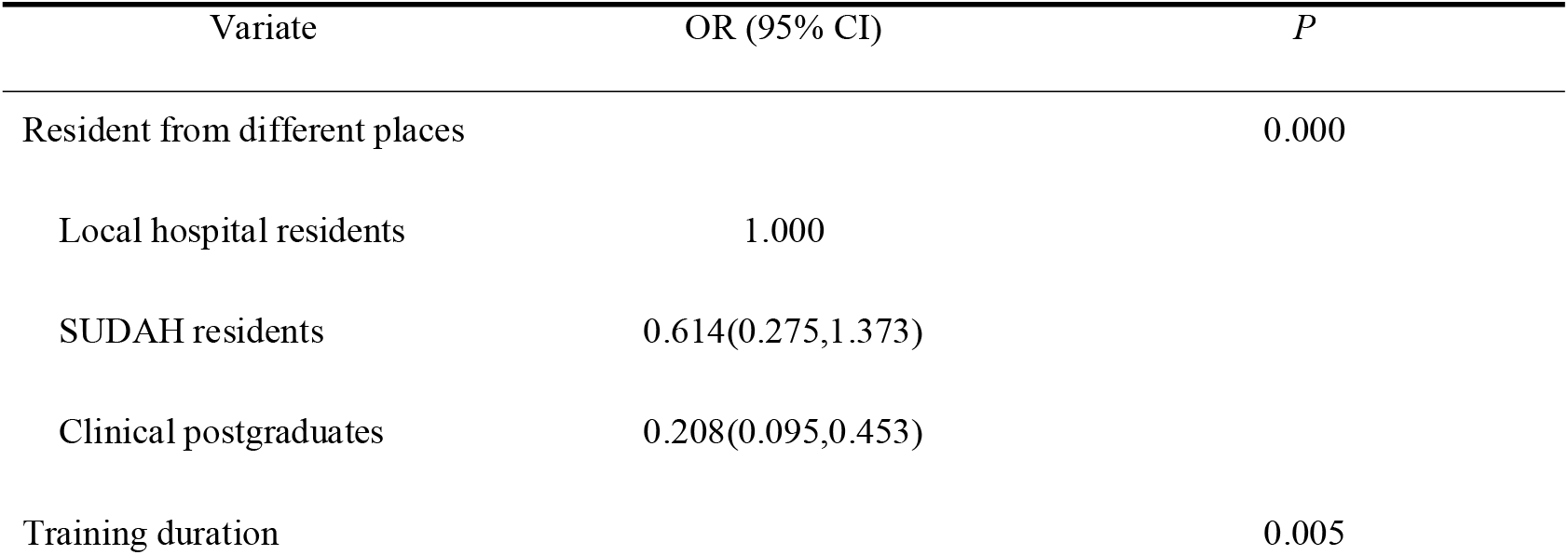
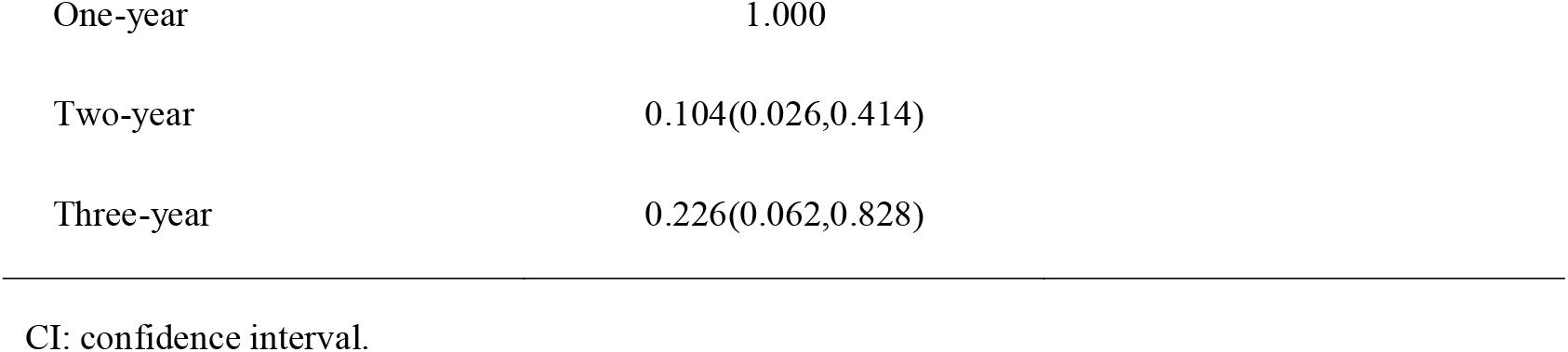
Predictors of qualification using multivariate Logistic regression analysis.

| Variate | OR (95% CI) | *P* |
| --- | --- | --- |
| Resident from different places |  | 0.000 |
| Local hospital residents | 1.000 |  |
| SUDAH residents | 0.614(0.275,1.373) |  |
| Clinical postgraduates | 0.208(0.095,0.453) |  |
| Training duration |  | 0.005 |

|  |  |
| --- | --- |
| One-year | 1.000 |
| Two-year | 0.104(0.026,0.414) |
| Three-year | 0.226(0.062,0.828) |

CI: confidence interval.

## Discussion

We found that the qualification rates of SRT programme differ significantly annually. And, age, gender, the highest medical education degree, residents from different places, training duration and years of medical school study were also different significantly year by year. The residents from local hospital and the short training duration were factors that affected the qualification rate of SRT programme of internal medicine.

Since the SRT programme gradually became compulsory medical education from 2014 [8], more and more medical graduates participated in this programme and thus enhanced their clinical competency. Because the present residency training programme integrated with the education of medical professional master’s degree, it received welcome from numerous undergraduates. Hence, our study showed that the percentage of clinical postgraduates was increasing and presented a trend of younger age year by year. The residents with bachelor’s degree accounted for an increasing proportion from 2015 to 2018. As a result, the proportion of residents with three-year training duration was increasing year by year. It was in accordance with the demands of medical and health industry, nowadays. Before practicing independently, residents would be benefited from the standardization training and improved their thinking abilities, decision-making abilities and practical abilities. These younger residents with a higher education degrees and stronger abilities are becoming the main force of doctors in China. It is conducive to construct a high-quality medical health care system and implement graded medical care to suit the new demands on medical and health services resulted from industrialization, urbanization, ecological environment changes as well as aging population. That is exactly the original intention of the reform of SRT programme launched by National Health Commission.

The “standardization” of SRT means to achieve homogeneity of residency training regardless the discrepancy among individuals and training bases. However, our study still showed some significant differences between the qualified group and the non-qualified group. We found that the residents in the non-qualified group were a little older than the qualified group. The proportion of residents with doctoral education degrees was higher in the non-qualified group than the other group, as well as the proportion of residents with one-year training duration. Compared with the qualified group, the residents in the non-qualified group had a few more years of medical school study and accounted for a higher proportion of whom ever worked in medical institutions before. All of these differences could be explained by the relative high proportion of doctoral degree in the non-qualified group. Most of these residents worked in laboratories during their education of doctoral degree. Due to lack of experience of clinical work, these residents could not pass the exams after only one-year training duration. Thereby, it is important to enforce the residency training for all medical graduates and maybe it was no matter their medical education degree. And, the longer training duration, the higher qualification rate of the SRT programme would be achieved.

And, the two groups of residents had significant discrepancy in the index of residents from different places, namely SUDAH residents, clinical postgraduates and local hospital residents. As we know, due to shortage of human resource for medical care, some low-resourced hospitals would sign labor contracts with medical graduates in advance. Then, those local hospital residents were sent for SRT program as “Unit Persons”, which was different with Shanghai municipal SRT policies [9]. The boldest aspect of the Shanghai pilot was designation of resident trainees as “Society Persons” rather than “Unit Persons” [3]. Society Persons referred to the residents who signed lab contracts with the Municipal health bureau rather than with specific hospital, which was similar with residents in the United Kingdom and the United States of America. They would look for steady jobs after qualification from SRT programme. Obviously, the job uncertainty added occupational stress together would make he or she be more aware of the programme’s importance and thus more supportive of its policies [10]. The job stability of Unit Persons had a shortcoming which posed a barrier to stimulate enthusiasm for learning and working in a certain measure. After transition from Unit Person to Society Person, it bought inherent challenges such as employment instability and less attachment to the training bases. The press would turn into their motivation for improvement in residents’ competency [11], while it might be associated with residency burnout [12–14]. Meanwhile, increasing residents’ awareness of the programme importance in advance during the residency training might reduce their stress and anxiety about the future, thus enhancing their professional competency [15,16]. Therefore, the actions to accelerate the transition from Unit Persons to Society Persons in the rest of China except Shanghai are urgent and will contribute to the ultimate “homogeneity” of SRT.

Although China has studied international experiences first, SRT as an innovative reform policy is still necessary to sum up experiences timely in the implementation process. We continuously adjust the detailed actions to meet the aims of SRT. Studies on SRT are still limited, especially on the evaluation of its effects. The present study analyzed the demographic and context data that could affect the performances of residents while ignoring factors related to burnout. A few studies on burnout indicated that it was universally existed in residency training [12,17,18] and might affect the residents’ competency. It prompts us to pay attention on this issue and make some adjustments. In future, we might deeply research the effects of burnout on the outcome of SRT.

## Supporting information

Figure 1. illustrate the gender distributions by the year

Figure 2. showed the percentages of training duration of three years increasing annually

Figure 3. illustrate the comparisons of each examination score annually

Figure 4. illustrated the differences of three types of identities of residents through the method of multiple comparisons of rate

## Acknowledgements

We thank all the staff from the Education Training Center for their kindly support during this work.

## Declaration of interest

The authors report no conflict of interest.

**LEGENDS FOR FIGURES: (supplements)**

Figure 1. illustrate the gender distributions by the year.

Figure 2. showed the percentages of training duration of three years increasing annually.

Figure 3. illustrate the comparisons of each examination score annually.

## Notes

**Source of funding support:** Supported by Jiangsu Province Hospital Association (No. JSYGY-3-2019-155).

### Competing Interest Statement

The authors have declared no competing interest.

