## Supplementary figures and images for "Factors that affected the qualification of standardized residency training of internal medicine: a prospective longitudinal study"

### Figure 1. illustrate the gender distributions by the year

Figure 1. illustrate the gender distributions by the year.

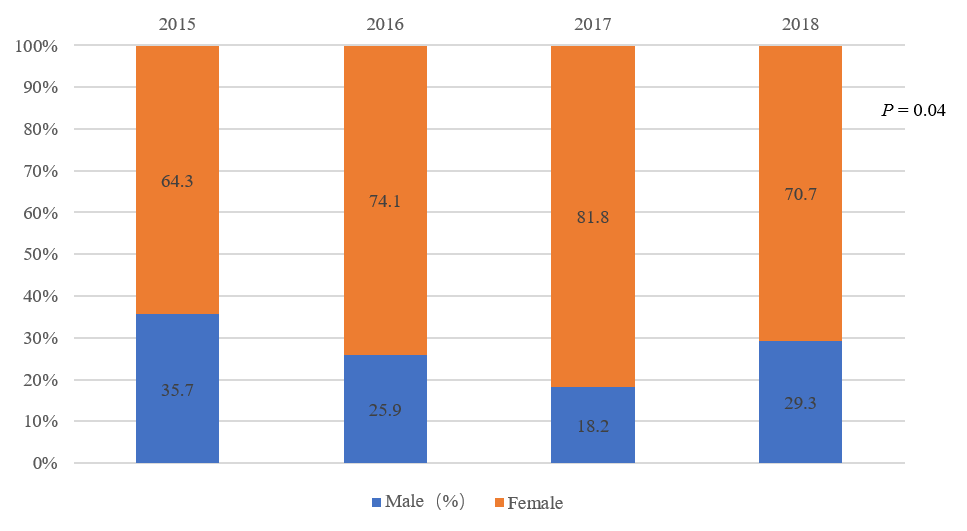

### Figure 2. showed the percentages of training duration of three years increasing annually

Figure 2. showed the percentages of training duration of three years increasing annually.

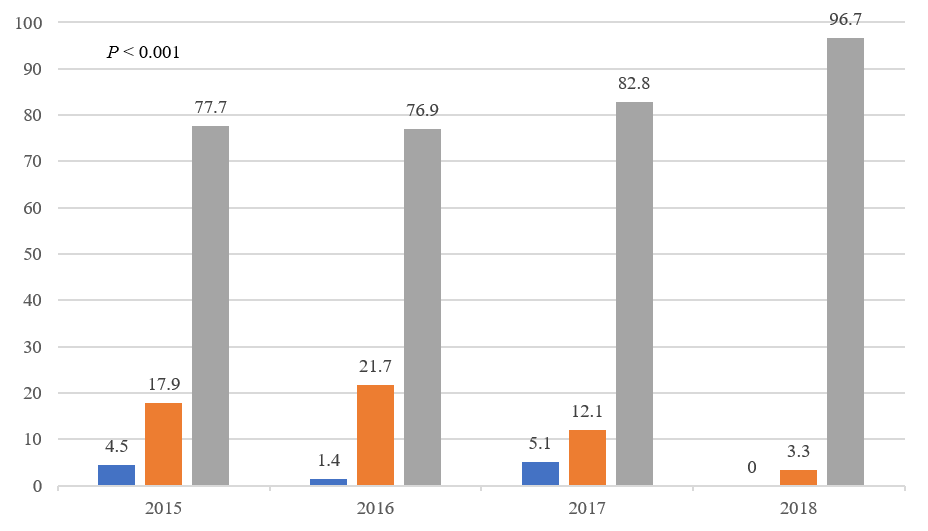

### Figure 3. illustrate the comparisons of each examination score annually

Figure 3. illustrate the comparisons of each examination score annually.

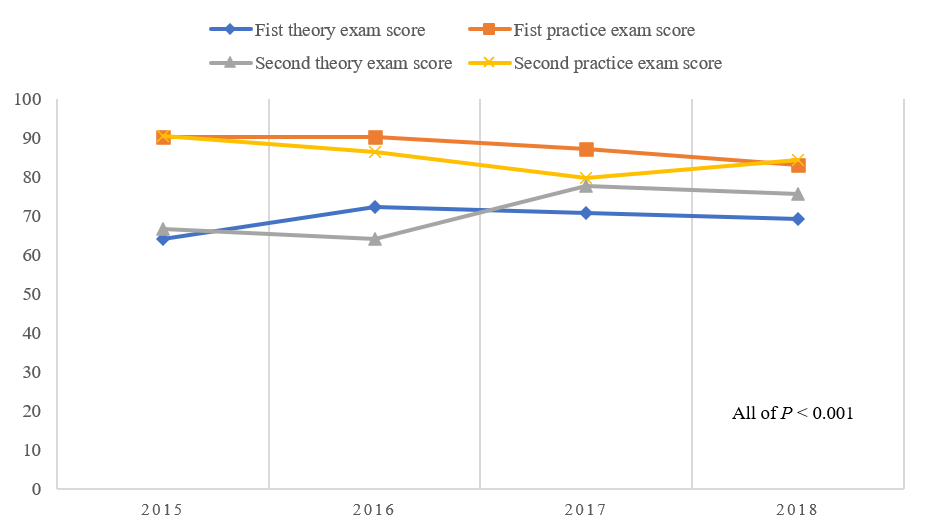
