## Supplementary material for "Factors that affected the qualification of standardized residency training of internal medicine: a prospective longitudinal study": Figure 4. illustrated the differences of three types of identities of residents through the method of multiple comparisons of rate

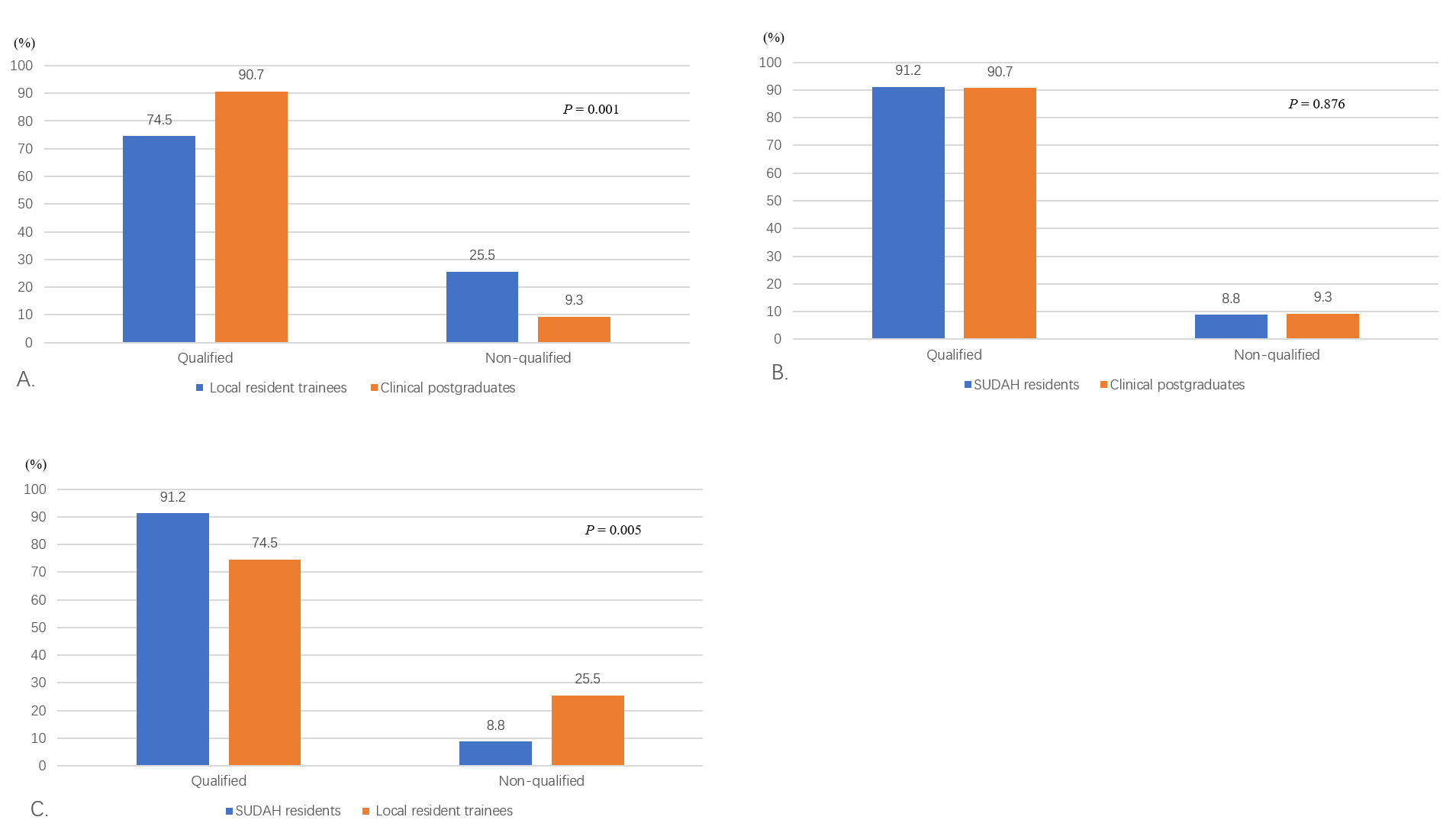
